# Androgen receptor activity inversely correlates with immune cell infiltration and immunotherapy response across multiple cancer lineages

**DOI:** 10.1101/2024.05.08.593181

**Authors:** Ya-Mei Hu, Faming Zhao, Julie N. Graff, Canping Chen, Xiyue Zhao, George V. Thomas, Hui Wu, Adel Kardosh, Gordon B. Mills, Joshi J. Alumkal, Amy E. Moran, Zheng Xia

## Abstract

There is now increasing recognition of the important role of androgen receptor (AR) in modulating immune function. To gain a comprehensive understanding of the effects of AR activity on cancer immunity, we employed a computational approach to profile AR activity in 33 human tumor types using RNA-Seq datasets from The Cancer Genome Atlas. Our pan-cancer analysis revealed that the genes most negatively correlated with AR activity across cancers are involved in active immune system processes. Importantly, we observed a significant negative correlation between AR activity and IFNγ pathway activity at the pan-cancer level. Indeed, using a matched biopsy dataset from subjects with prostate cancer before and after AR-targeted treatment, we verified that inhibiting AR enriches immune cell abundances and is associated with higher IFNγ pathway activity. Furthermore, by analyzing immunotherapy datasets in multiple cancers, our results demonstrate that low AR activity was significantly associated with a favorable response to immunotherapy. Together, our data provide a comprehensive assessment of the relationship between AR signaling and tumor immunity.

## Introduction

The androgen receptor (AR) is a nuclear transcription factor (TF) belonging to the steroid hormone receptor superfamily^1^. The human AR protein, primarily responsible for mediating the biological actions of androgens (male sex hormones), is composed of 919 amino acids and is encoded by a 180 kb AR gene located on chromosome Xq11-12^2, 3, 4^. Similar to other steroid receptors, AR consists of three main functional domains: the N-terminal transcriptional regulation domain (NTD), the DNA-binding domain (DBD), and the ligand-binding domain (LBD)^5^. In the canonical AR signaling pathway, AR is activated by binding to its native ligands, dihydrotestosterone (DHT) or testosterone, leading to conformational changes, homodimer formation, and nuclear translocation^3, 6^. The nuclear AR then binds to androgen response elements (AREs) of target genes for transcriptional regulation, crucial for cell differentiation and proliferation^7, 8, 9^. Activation of AR signaling is essential for reproductive development in both sexes^10, 11^. AR is present in various cells and tissues, including the prostate, bone, muscle, adipose tissue, immune, reproductive, cardiovascular, neural, and hematopoietic systems, and has significant physiological effects even in non-reproductive tissues^12^.

Androgens/AR signaling is widely recognized as a critical factor in the development and progression of prostate and breast cancer^13, 14^. Cancer is a complex disease where malignant cells interact with the tumor microenvironment (TME), which includes immune cells, mesenchymal-origin cells, and the extracellular matrix^15^. The immunological component of the TME can either suppress or promote tumor development, with infiltrating immune cells playing a crucial role in influencing tumor growth, progression, therapeutic outcomes, and patient prognosis^15^. Several studies utilizing conditional AR knockout mice have demonstrated that androgens/AR are involved in immunomodulation^16, 17^. AR expression has been detected in almost all immune cells, and many immune-related genes have AREs in their promoters, thereby influencing both innate and adaptive immunity^16, 18^. For example, androgens/AR can affect cytokine production from various immune cells, regulate macrophage recruitment^19^, differentiation, and polarization^20^, and exert suppressive effects on the development and activation of T and B cells^21^. Moreover, recent evidence suggests that AR signaling plays a vital role in anticancer immunity not only in reproductive cancers^22^ but also in non-reproductive cancers^23, 24, 25^. Guan et al. reported that AR represses *Ifng* transcription in T cells, and AR inhibition with enzalutamide restored IFNγ production, which improved responsiveness to programmed death-1 (PD-1) targeted therapy in metastatic castration-resistant prostate cancer (mCRPC)^22^. Studies using preclinical models of bladder cancer^24, 25^ indicate that T cell-intrinsic AR signaling promotes the programmed exhaustion of CD8^+^ T cells in males, which may contribute in part to sex differences in cancer incidence and severity. There is mounting evidence to suggest that AR signaling plays a role in promoting cancer growth and progression across an array of malignancies, such as bladder, kidney, pancreatic, liver, endometrial, mantle cell lymphoma, salivary gland cancers, and others^13^. However, the role of AR activity in cancer immunity requires further examination.

In this study, we conducted a pan-cancer analysis of AR activity across 33 cancers using datasets from the Cancer Genome Atlas (TCGA). We defined the AR activity across cancer types, including its association with patient survival, specific gene expression programs, and correlation with immune cell abundance. We also explored the association between AR activity and immune signatures of interest. Subsequently, we investigated AR activity and immune cell profiles in patients with mCRPC and compared baseline and AR inhibitor-treated samples from the same patients. Furthermore, we unveiled an association between AR activity and tumor immunotherapy response using six independent datasets. Our study provides evidence for the potential importance of AR activity as a predictive biomarker of immunotherapy response in different malignancies.

## Results

### AR activity profiles in human cancers

To profile AR signaling activities in human malignancies, instead of using the expression levels of individual genes, we utilized a regulon-based analysis. We calculated AR activity using the Virtual Inference of Protein-activity by Enriched Regulon analysis (VIPER) analysis^26^, which employs the transcriptional regulatory network, including TFs and their targeted genes, to infer TF activity. Such a regulon-based method is more robust to noise and provides greater sensitivity in characterizing TF activities compared to measuring the gene expression of the TF itself^26^. **Figure 1a** depicts the AR activity across all TCGA tumors (10,340 samples in 33 TCGA studies). These 33 cancers exhibited significant differences in AR activity (ANOVA test, *p* < 2.2e−16), with each cancer type displaying a wide range of AR activity. Overall, prostate adenocarcinoma (PRAD) has the highest levels of AR activity on average compared to other tumors, while acute myeloid leukemia (LAML) has the lowest (**Fig. 1a**). TCGA study abbreviations and sample size for each dataset are summarized in Supplementary Table 1.

**Fig. 1.**
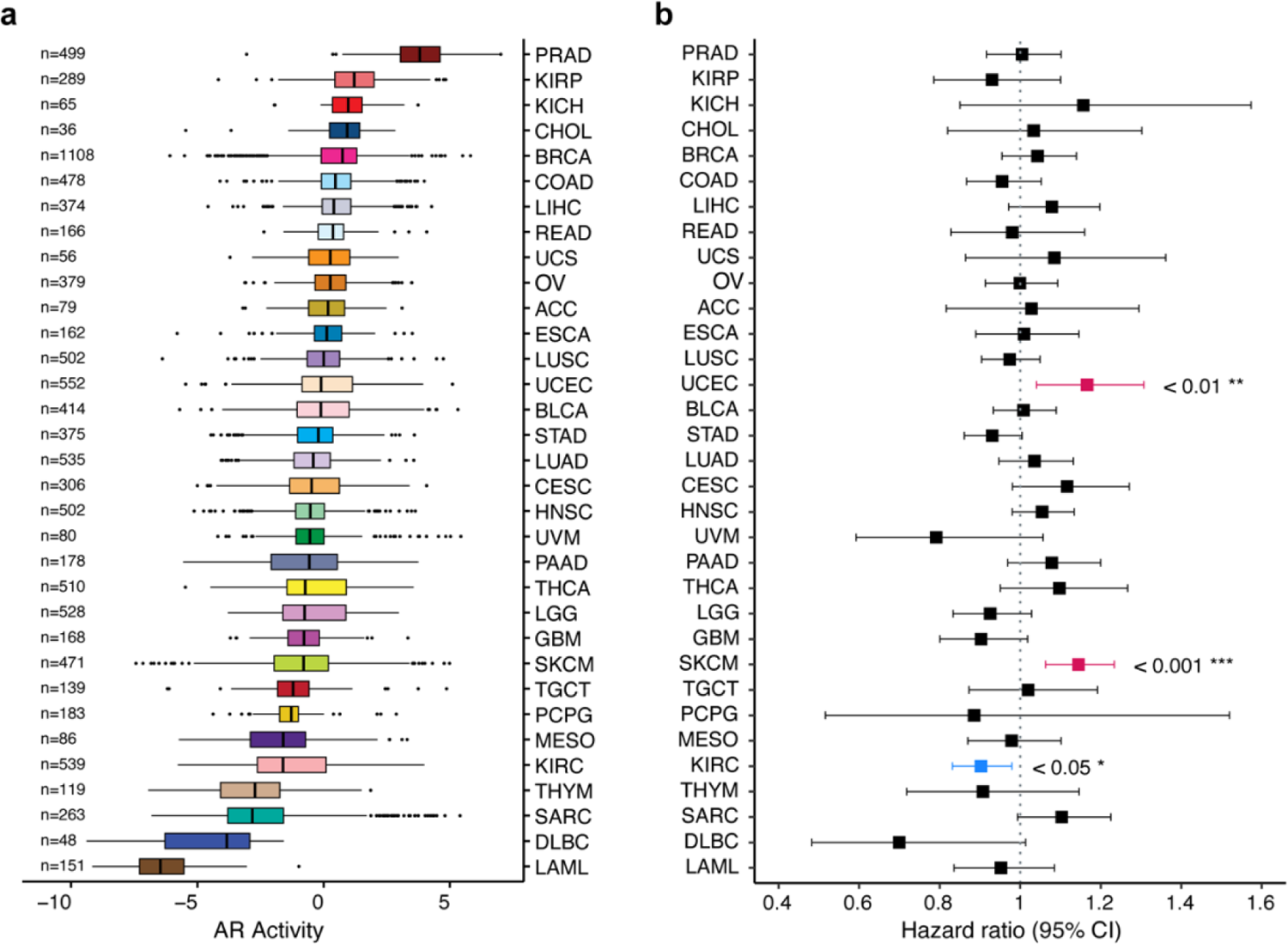
Overview of AR Activity and association with survival outcomes across 33 TCGA cohorts. **a**, Boxplots displaying AR activity sorted in order of decreasing AR activity median among 33 TCGA cancer types. Values on the left correspond to the total number of tumor samples (n) analyzed in each cancer cohort. TCGA study abbreviations and sample size for each dataset are listed in Supplementary Table 1. The center line indicates median, bounds of the box indicate upper and lower quartiles, whiskers indicate minimum and maximum, and outliers are marked with dots. **b,** Univariate Cox regression analysis of the AR activity on overall survival or progression-free interval (BRCA, DLBC, LGG, TGCT, THYM, PRAD, READ, THCA) endpoints according to Liu et al.^27^ recommendations was conducted in TCGA cohorts. Forest plots showing hazard ratio (HR) estimates, 95% confidence intervals (CI), and corresponding *p*-values. Cancers in which AR activity is significantly associated with good prognosis are highlighted in blue, and those significantly associated with poor prognosis are highlighted in hot pink color.

### The association between AR activity levels and patient survival outcomes

To gain insight into the clinical relevance of AR activity at the pan-cancer level, we conducted survival analysis in the 33 TCGA cohorts. According to the clinical endpoint usage recommendations for each cancer type^27^, progression-free interval (PFI) endpoints were used for survival analyses in 8 cancer types (breast invasive carcinoma [BRCA], lymphoid neoplasm diffuse large B-cell lymphoma [DLBC], brain lower grade glioma [LGG], testicular germ cell tumors [TGCT]), Thymoma [THYM]), PRAD, rectum adenocarcinoma [READ]), and thyroid carcinoma [THCA]); For the remaining cancer types, overall survival (OS) data were utilized.

The result indicated that high AR activity is associated with a decreased risk for kidney renal clear cell carcinoma (KIRC) (HR = 0.9, 95% CI 0.83–0.98, *p* = 0.014; **Fig. 1b**), and an increased risk for uterine corpus endometrial carcinoma (UCEC) (HR = 1.16, 95% CI 1.04–1.31, *p* = 8.2e-03; **Fig. 1b**) as well as for skin cutaneous melanoma (SKCM) (HR = 1.14, 95% CI 1.06–1.23, *p* = 3.6e-04; **Fig. 1b**). The remaining cancer types did not exhibit significant differences in the relationship between AR activity and survival outcomes of cancer patients.

### AR activity-correlated genes are significantly associated with immune system processes

We then investigated the potential impact of AR activity on other gene expression programs. To do so, we calculated Pearson correlation coefficients and *p*-values to assess the relationship between AR activities and the expression of each gene in each cancer type. The genes that are significantly correlated with AR activity were identified with *p* < 0.05. To determine the prevalence of AR activity-correlated genes across different tumors, we identified 31 genes significantly associated with AR activity across all tumor types (**Fig. 2a**). Remarkably, 99.7% of the correlations between the 31 genes and AR activity were negative (1020 negative correlation coefficients, 3 positive correlation coefficients), with 632 of 1023 coefficients being strongly negative (less than −0.5). Notably, several MHC class I and II genes (HLA-E, HLA-F, HLA-DPB1, HLA-DRB1 and HLA-DMA), as well as T-cell and B-cell specific genes (CD2, CD6, CD7, CD247 and CD72) are among the 31 genes most frequently and negatively correlated with AR activity (**Fig. 2a**). Additionally, we used Gene Ontology (GO) biological process and Reactome pathway analysis to gain further insight into the underlying biology of these 31 genes. These genes were predominantly involved in immune processes such as T cell activation regulation, leukocyte activation, leukocyte cell-cell adhesion, T cell proliferation, antigen processing and presentation, TCR signaling, and interferon signaling (**Fig. 2b, c**). Such unbiased analyses revealed that AR activity is negatively correlated with genes intensively involved in positive immune processes, strongly suggesting immunosuppressive effects of AR in cancer patients.

**Fig. 2.**
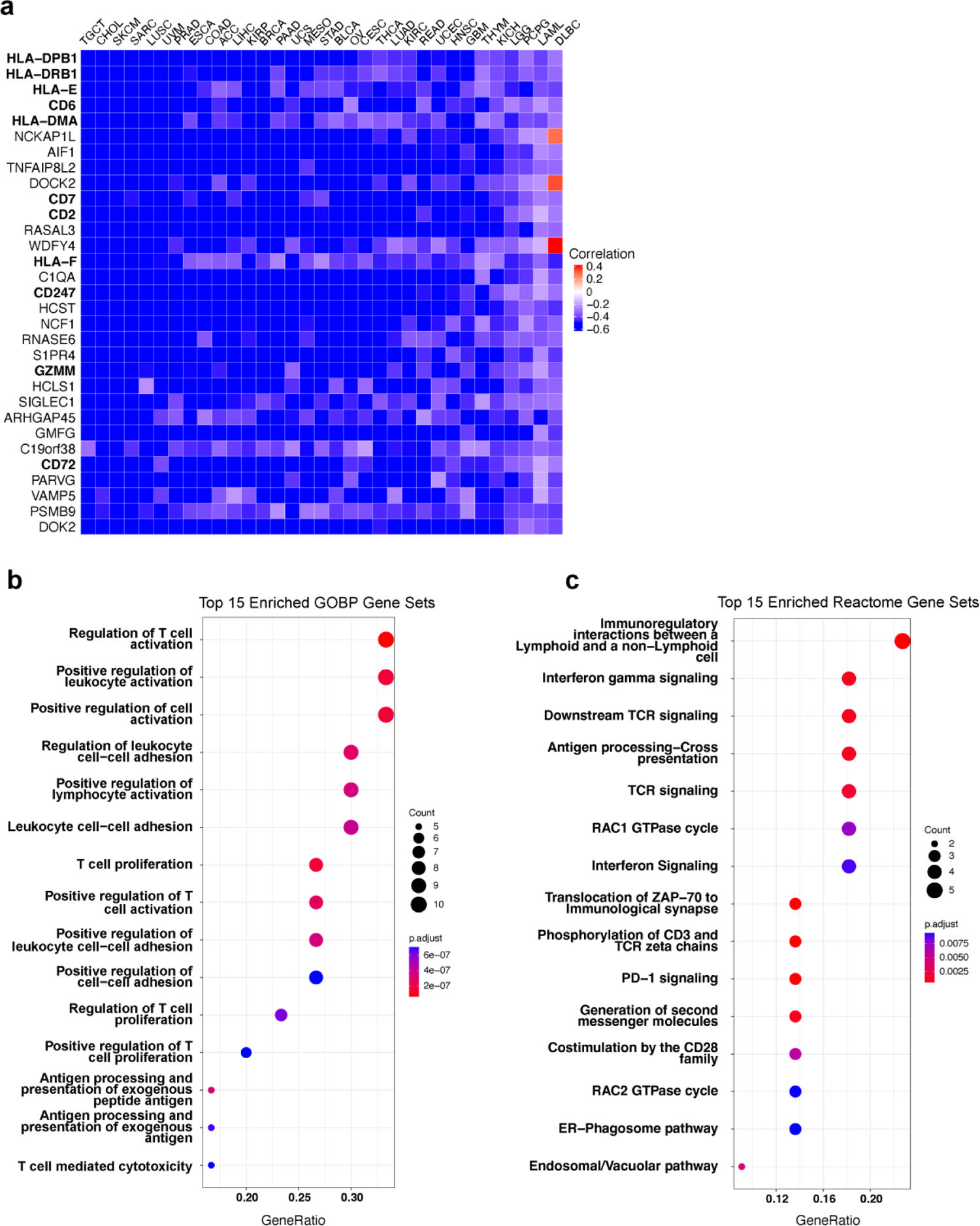
AR activity shows a strong negative correlation with genes involved in immune system processes. **a**, Evaluation of correlation between AR activity and each gene expression within the entire dataset of each tumor type across all 33 TCGA cohorts. Heatmap displaying the 31 genes (rows) that are significantly correlated with AR activity across all 33 tumor types. Pearson correlation tests were used to calculate the correlation between AR activity and each gene’s expression in each cancer type. Significant correlations (*p* < 0.05) are indicated in red (positive correlation) or blue (negative correlation). Cancer types (columns) are sorted in order of decreasing mean of correlation coefficient. **b, c** Pathway enrichment analysis of the 31 AR activity-correlated genes. Dot plots indicating the top 15 significantly enriched Go BP terms (b) and Reactome pathways (c). The vertical axes represent the function classifications or pathways, and the horizontal axes represent the ratio of genes contained in the ontology/pathways. The dot’s color indicates the adjusted *p*-value of enrichment significance, and the size of the dot indicates the number of genes in the functional class or pathway. GO, Gene Ontology; BP, biological process.

### AR activity negatively correlates with immune infiltration in cancer

Because of our observation that AR activity is negatively associated with expression levels of immune-related genes, we next investigated the correlation between AR activity and immune cell composition in tumors by deconvoluting TCGA bulk RNA-seq samples. Similarly, we found that AR activity negatively correlates with most immune infiltrates estimated by TIMER analysis of bulk RNA-seq, including B cells, CD4^+^ T cells, CD8^+^ T cells, neutrophils, macrophages, and myeloid dendritic cells (**Fig. 3a**). Among 33 TCGA cancer types, AR activity shows significant moderate to strong negative correlations (correlation coefficients ranging from −0.4 to −0.8) with the abundance of six immune cell types in 19 cancer types as estimated by TIMER. Particularly, Sarcoma (SARC) and SKCM exhibited the highest degree of negative correlations between AR activity and immune infiltration (**Fig. 3a, b**). Beyond the 6 immune cell types estimated by TIMER, we also conducted single-sample gene set enrichment analysis (ssGSEA) to quantify the degree of infiltration of 28 human immune cell categories^28^ in each bulk RNA-seq sample, and computed its correlation with AR activity (Supplementary Fig. 1). Once again, AR activity showed a significantly negative correlation with immune cell infiltration across cancer lineages. Moreover, recognizing sex as a biological variable that influences various immune system functions, we assessed differences in immune infiltration between male and female patients (**Fig. 3c**). Male patients with SARC showed higher infiltrating levels of CD8^+^ T cells, neutrophils, macrophages, and myeloid dendritic cells while female patients with LUSC and lung adenocarcinoma (LUAD) have higher infiltration levels of multiple immune cell types (**Fig. 3c**).

**Fig. 3.**
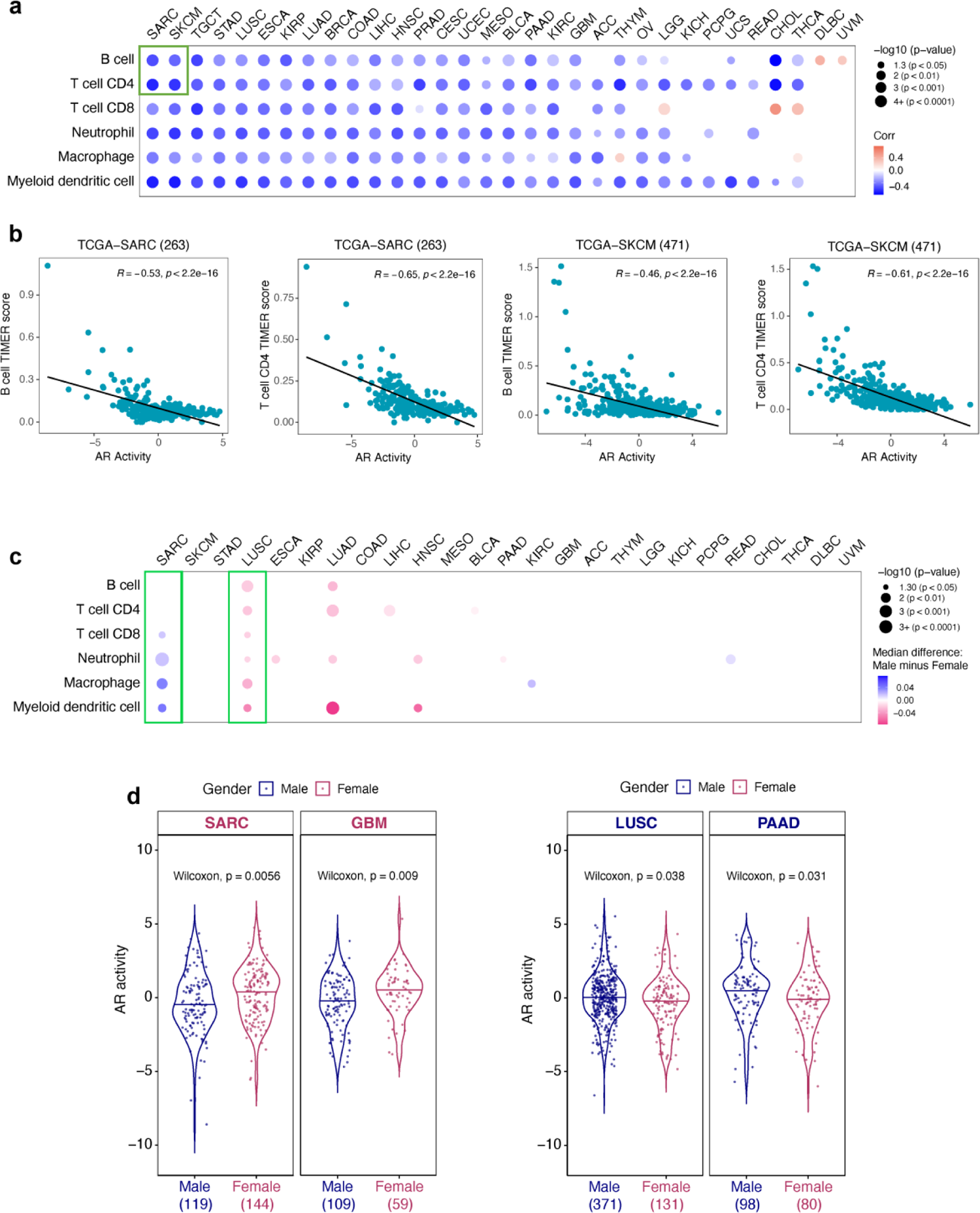
AR activity exhibits a negative correlation with immune infiltration in cancers. **a** Correlations between AR activity and six immune cell populations (B cells, CD4^+^ T cells, CD8^+^ T cells, macrophages, dendritic cells, and neutrophils) across 32 cancer types (LAML is not applicable in TIMER calculation). Each circle represents a correlation coefficient value analyzed by two-tailed Pearson correlation test. Positive correlation coefficients are displayed in orange and negative correlation coefficients in blue color. The color intensity is proportional to the correlation coefficients. The circle size corresponds to the *p*-value, with correlation coefficients having *p*-value > 0.05 left blank. Cancer types (columns) are sorted in order of decreasing mean of correlation coefficient. The enrichment scores of immune infiltration levels are determined by TIMER algorithm. **b**, Representative scatter plots demonstrating the dots in the green square of Fig. 3a that show the correlation between AR activity and B cell or CD4^+^ T cell in SARC and SKCM cohorts. **c,** Differences in tumor immune infiltration levels between male and female patients in 25 TCGA cohorts. The dots indicate significant median differences in tumor immune infiltration levels, with *p*-values > 0.05 left blank. Blue and pink dots represent higher immune infiltration levels in males and females, respectively. The color intensity is proportional to the number of median differences. The circle size corresponds to the *p*-value calculated using a two-sided Wilcoxon rank-sum test. Cancer types (columns) are sorted in accordance with Fig. 3a. **d**, Violin plots illustrating AR activity differences in four TCGA cohorts between male (navy) and female (maroon) patients. Statistical significance was determined using two-sided Wilcoxon rank-sum tests between males and females. The values at the bottom indicating the total number of tumor samples in each condition.

Meanwhile, we also investigated the differences in AR activity between sexes for 27 cancer types, each with at least 12 samples from both female and male tumors (**Fig. 3d**, Supplementary Fig. 2). AR activity was higher in male tumors in lung squamous cell carcinoma (LUSC) and pancreatic adenocarcinoma (PAAD), whereas AR activity was higher in female tumors in brain lower grade glioma (GBM) and sarcoma (SARC) (**Fig. 3d**, *p* < 0.05). Notably, female patients with SARC and male patients with LUSC exhibited higher AR activity (**Fig. 3d**) but lower immune infiltrates (**Fig. 3c**), suggesting AR signaling activation may contribute to reduced immune infiltration observation in these populations (**Fig. 3c**). We further evaluated the association between AR activity and immune cell abundances in male and female patients, respectively (Supplementary Fig. 3). Finally, considering the impact of steroid hormones on immunity and their implication in various cancers, we also assessed the correlation between the activity of estrogen receptors, progesterone receptor, and glucocorticoid receptor with immune infiltrates. The results revealed predominantly positive correlations across most cancer types, except for THYM, which showed moderate to strong negative correlations (Supplementary Fig. 4). Taken together, AR activity was negatively correlated with immune infiltration in both males and females. The results from multiple angles implied that AR signaling activation participates in the immune infiltration process.

### AR activity negatively correlates with prognostic immune signatures in cancer

To further examine whether AR activity is associated with immunological activity in TME, we investigated three IFN-γ-related immune gene expression signatures and one signature of tertiary lymphoid structures (TLSs) known to be associated with favorable responses to immune checkpoint inhibitors and of prognostic value. These included the Hallmark interferon-gamma signaling (IFN-γ)^29^, the 18-gene T cell-inflamed gene expression profile (GEP) ^30^, the immunologic constant of rejection (ICR)-20 genes signature^31^, and the TLS signature (Cabrita et al.^32^). We performed ssGSEA to calculate immune signature scores in the TCGA bulk RNA-seq dataset and analyzed the correlation between AR activity and the multigene immune signature scores. Significant negative correlations were observed between each signature score and AR activity in all TCGA tumor samples (n=10,340), with each having a Pearson correlation value ranging from −0.39 to −0.44 (**Fig. 4a-d**). The correlations between AR activity and each immune signature score within each cancer type are displayed in **Figure 4e-h**. In most cancers, except DLBC and THYM, robust and significant inverse relationships concordantly existed between AR activity and the prognostic immune signatures. Strong negative correlations were observed in around one-third of cancers (Pearson correlation < −0.6, *p* < 2.2e-16). These results suggested that AR signaling is highly involved in tumor immunity across cancers. Furthermore, we also explored the correlation between AR activity and the signature activities in normal tissue from The Genotype-Tissue Expression (GTEx) database^33^ (**Fig. 5**, Supplementary Fig. 5, Supplementary Fig. 6, Supplementary Table 2). Once again, AR activity exhibited significant negative correlations with these immune signature activities in GTEx samples, indicating AR signaling may have immunosuppressive effects in both normal and tumor tissues.

**Fig. 4.**
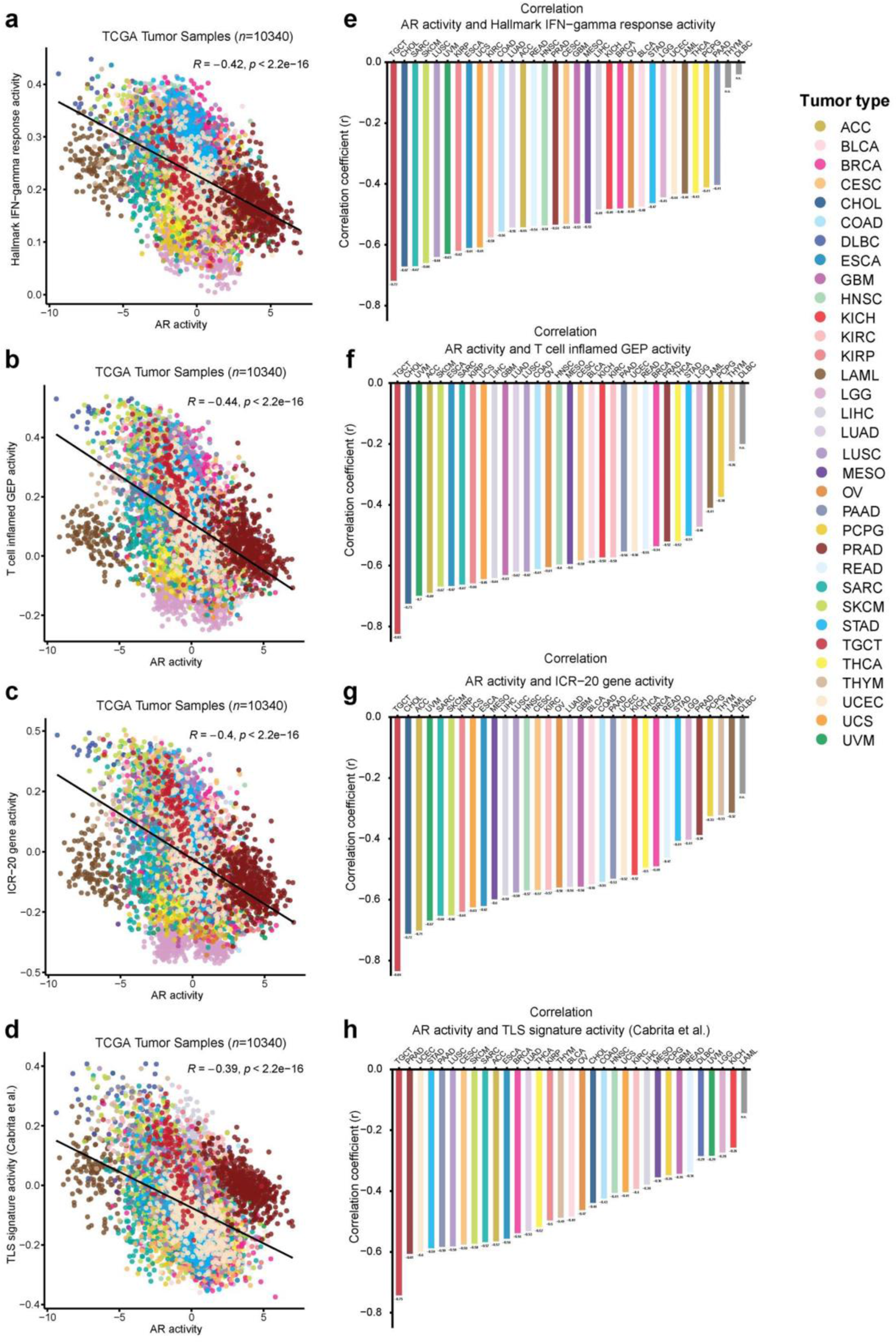
AR activity levels are negatively correlated with gene signatures of three prognostic gene signatures and TLS within 33 TCGA cohorts. Scatter plots (**a, b, c, d**) showing the Pearson correlation of AR activity with **a** Hallmark IFN-γ pathway, **b** T cell–inflamed GEP, **c** ICR-20 gene, and **d** TLS signature activity scores of all TCGA tumor samples. Each dot represents one tumor sample (n=10,340), with colors indicating different tumor types. Tumor types are listed on the right by color code (n = 33). Bar plots (**e, f, g, h**) displaying the Pearson correlation of AR and signature activities within each cancer type. The X-axis denotes cancer types arranged in decreasing correlation coefficient value (Y-axis). The texts below each bar indicate the Pearson correlation coefficients. Bars on the far right with a *p*-value > 0.05 are colored in light grey. GEP: gene expression profile. ICR: immunologic constant of rejection. TLS: tertiary lymphoid structures. ns: non-significant.

**Fig. 5.**
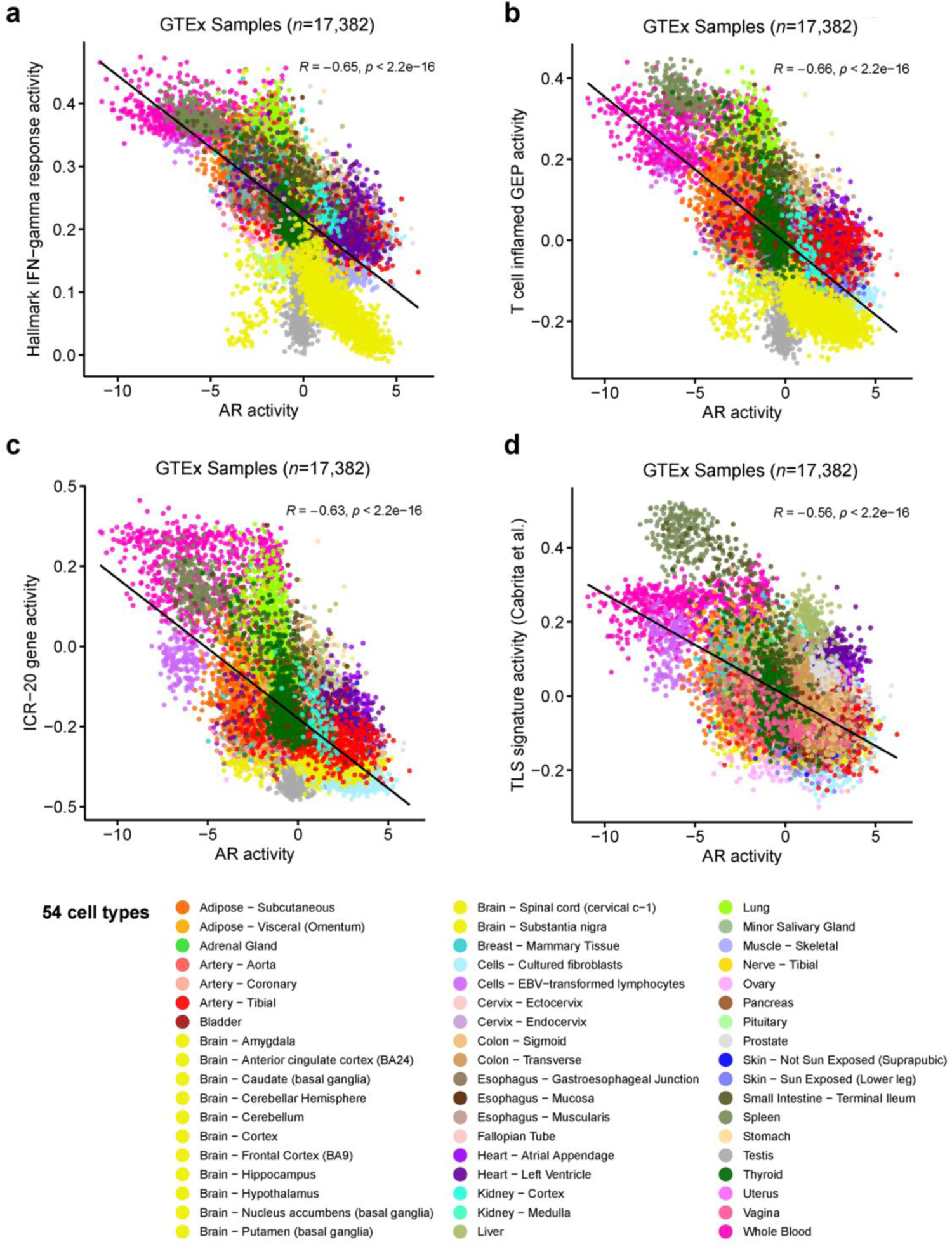
AR activity levels are negatively correlated with gene signatures of IFN-γ signaling and TLS in GTEx tissue samples. Scatter plots (**a, b, c, d**) showing the Pearson correlation of AR activity with **a** Hallmark IFN-γ pathway, **b** T cell–inflamed GEP, **c** ICR 20-gene, and **d** TLS signature activity scores of all GTEx tissue samples. Each dot represents one normal tissue sample (*n*=17,382), with colors indicating 54 different tissue types. Tissue types are listed in the legend by color code. The AR activity profiling among all tissue types in the GTEx database is presented in Supplementary Fig. 5. The Pearson correlation of AR activity and signature activity scores within each tissue type is presented in Supplementary Fig. 6. GEP: gene expression profile. ICR: immunologic constant of rejection. TLS: tertiary lymphoid structures. GTEx: The Genotype-Tissue Expression.

### Inhibiting AR signaling increases immune infiltration and IFN-γ signaling activity in mCRPC patients

Based on these findings, we hypothesized that AR-targeting treatments would impact the immune profiles of patients. We conducted RNA expression profiling of tumor tissues from matched patient biopsies before and after treatment with the AR signaling inhibitor enzalutamide (Enza), one of the principal treatments for mCRPC^34^. (**Fig. 6a**). Indeed, master regulator analysis by comparing the two conditions identified that AR activity was repressed in tumor samples post-treatment (**Fig. 6b**), confirming that Enza effectively inhibited AR activity in these patients. Consequently, functional signature enrichment analyses showed that, following treatment, tumor samples had increased levels of immune cell abundances in 19 out of 28 immune cell types (**Fig. 6c**), and exhibited significant upregulation of genes involved in multiple positive immune process pathways in GO BP terms (**Fig. 6d**). Moreover, GSEA analysis demonstrated significant enrichments of prognostic immune signatures in tumor samples after AR signaling inhibition by Enza treatment, including the IFN-γ signaling signature, the 18-gene T cell-inflamed GEP and ICR 20-gene signatures (**Fig. 6e**). These observations suggest that AR inhibition leads to increased immune cell infiltration and could potentially enhance the response to immune checkpoint blockade (ICB).

**Fig. 6.**
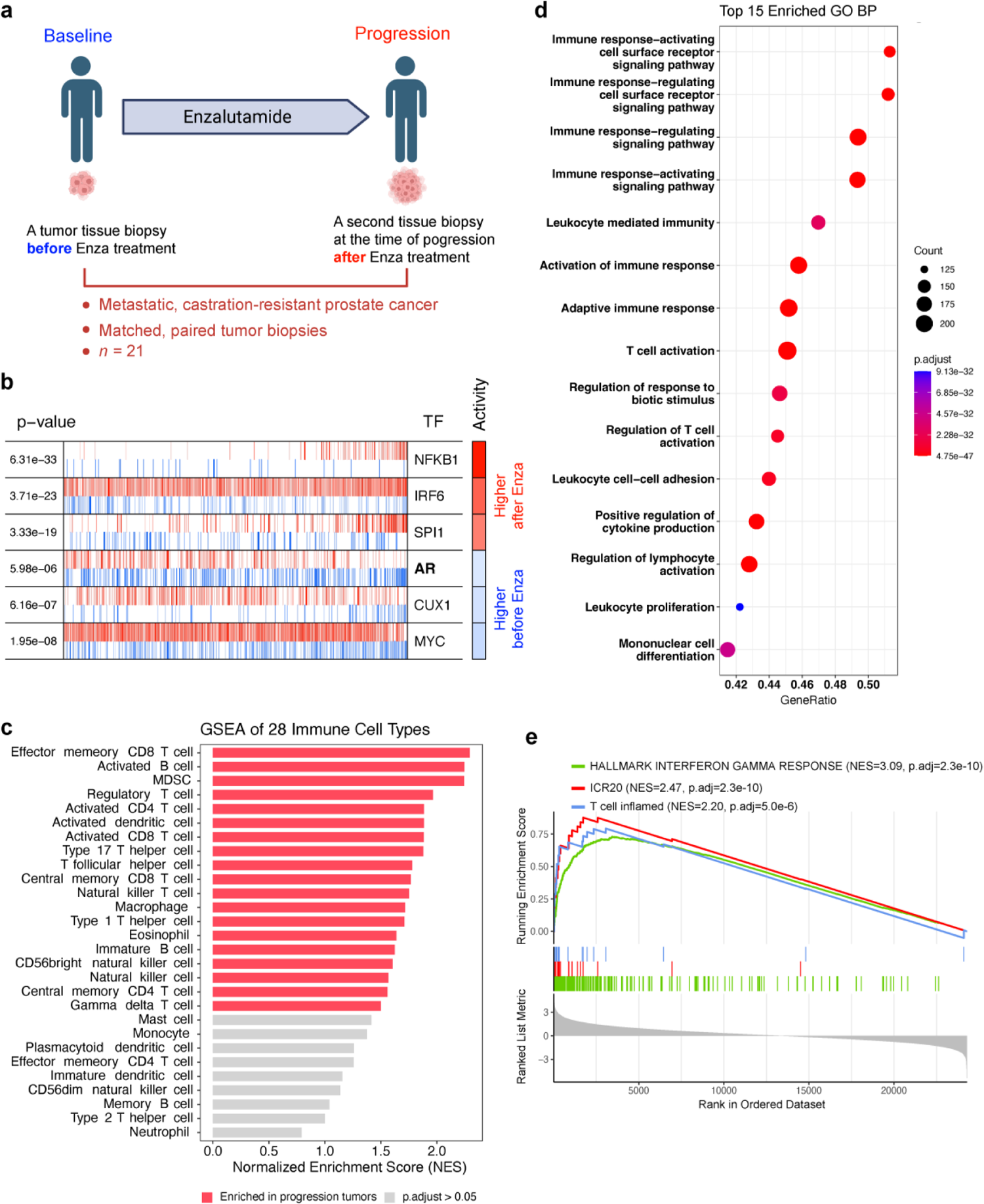
Master regulator and pathway enrichment analysis revealed decreased AR activity, increased immune cell abundance, and the enrichment of interferon-gamma response after enzalutamide treatment in the 21 matched biopsy mCRPC patients. **a**, Study schematic. Created with BioRender.com. **b**, msVIPER plot depicting the top three TFs predicted to be most activated (red in Activity) or deactivated (blue) in progression tumor samples compared with baseline tumor samples. Tick marks in red or blue lines represent targets of TFs that are positively or negatively regulated, respectively. **c**, GSEA of 28 immune cell types indicating immune infiltration in progression tumor samples. **d**, GSEA of GO biological process demonstrating the top 15 enriched pathways (with NES ranging from 2.71 to 2.82) in progression tumor samples associated with immune system processes. **e**, GSEA plot showing significantly activated three prognostic gene signatures after Enza treatment. CRPC: castration-resistant prostate cancer. TFs: transcription factors. GSEA: Gene Set Enrichment Analysis. GO: Gene Ontology. BP: biological process. ICR: immunologic constant of rejection.

### AR activity level inversely correlated with response to ICB treatment

Because the immunological activity within the TME contains prognostic information and because of the observed inverse relationships between AR activity and immune signatures, we hypothesized that AR activity levels in patient baseline tumors might correlate with their response to immunotherapy. To test this hypothesis, we first investigated the correlation between AR activity and the three prognostic immune signatures using six publicly available RNA-Seq datasets from pretreatment tumor samples that include response data after ICB treatment from various cancer (GSE145996, phs000452.v2.p1, GSE135222, GSE126044, Yang et al., and Guan et al., **Fig. 7**). Strong negative correlation (Pearson correlation < −0.5) was observed in almost all datasets between AR activity and the prognostic immune signatures (**Fig. 7a-p**). Subsequently, we compared AR activity levels between non-responders and responders in these cancer cohorts. We found that responders exhibited a significantly lower AR activity level than non-responders in melanoma, NSCLC, and mixed tumor cohorts, respectively (*p* < 0.05, two-sided Wilcoxon rank-sum test; **Fig. 7q-s**). Of note, responders had a trend toward lower AR activity in mCRPC patients (*p* = 0.13) (**Fig. 7t**). Our results indicated that AR signaling correlates strongly with cancer patients’ response to ICB. This suggests that the level of AR activity in baseline tumors may provide insight into pre-existing tumor immune microenvironment and influence therapy selection.

**Fig. 7.**
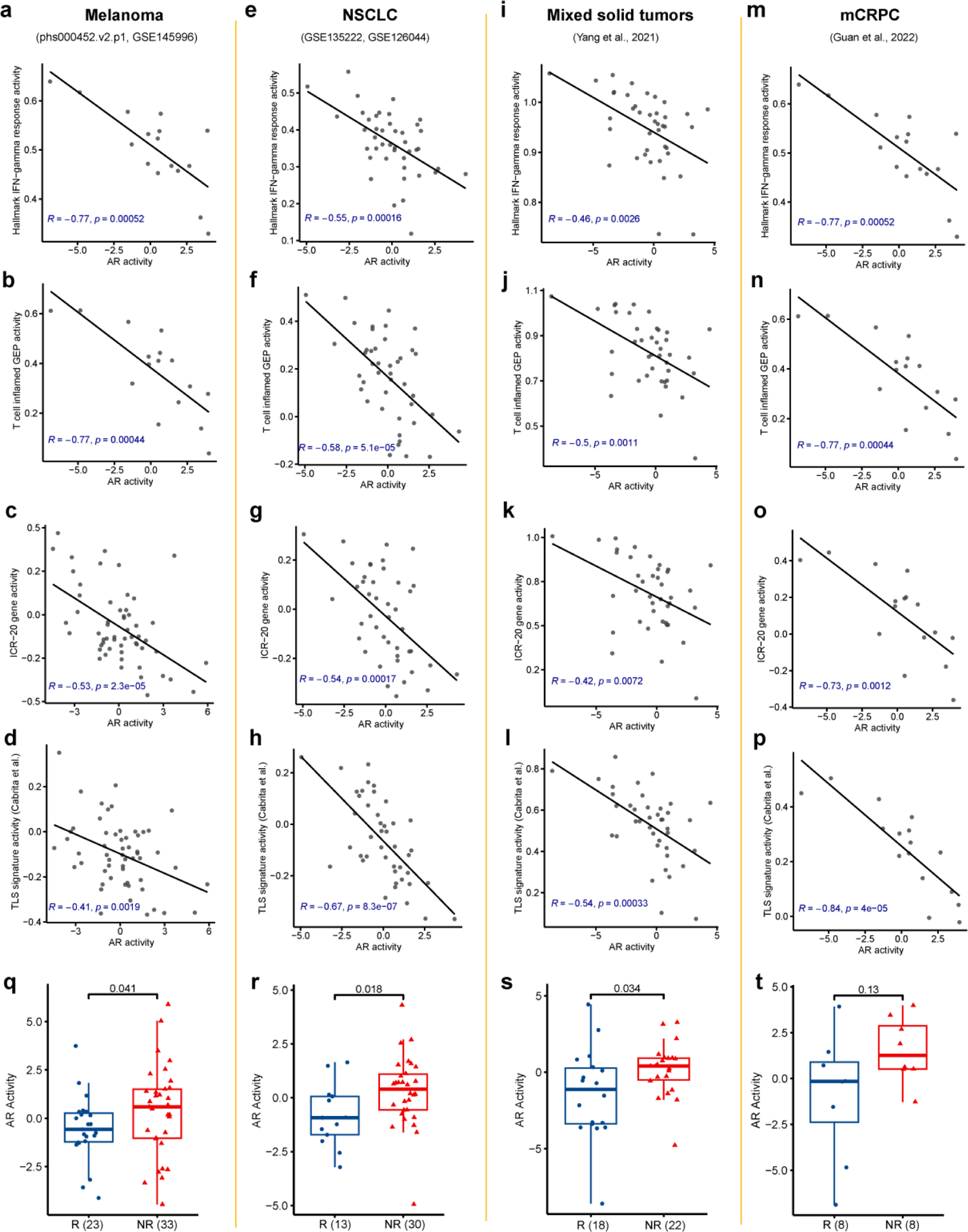
AR activity level inversely correlated with response to ICB treatment in public datasets. **a-l**, Scatter plots showing Pearson correlations of AR activity and the Hallmark IFN-γ pathway, T cell–inflamed GEP, ICR-20, and TLS gene signature activity scores for the phs000452.v2.p1 and GSE145996 combined melanoma dataset (**a-d**), the GSE135222 and GSE126044 combined NSCLC dataset (**e-h**), Yang et. al mixed tumors dataset (**i-l**), and the Guan et. al AR prostate cancer dataset (**m-p**). **q-t,** Boxplot displaying AR activities between responders (R) and non-responders (NR) to ICB treatment in the melanoma dataset (**m**, *p* = 0.041), the NSCLC dataset (**n**, *p* = 0.018), the mixed tumors dataset (*p* = 0.034), and Guan et. al AR prostate cancer dataset (**p**, *p* = 0.13). GEP: gene expression profile. ICR: immunologic constant of rejection. TLS: tertiary lymphoid structures. NSCLC: non-small-cell lung cancer. ICB: immune checkpoint blockade.

### AR activity is inversely correlated with immune infiltration from single-cell analysis

To further validate the relationship between tumor cell-intrinsic AR signaling and immune infiltration at a single-cell resolution, we initially inquired a comprehensive scRNA-seq atlas of human prostate cancer (PCa) from our recent study^35^ (**Fig. 8a, b**; tumor samples = 66, total cells = 210,879). The results showed a significant negative correlation between AR activity in the epithelial compartment and the level of immune cell infiltration in the TME of human PCa (**Fig. 8c**). Additionally, odds ratio (OR) analysis^36^ revealed that most immune cell types were significantly enriched in ARlow tumor tissues (OR > 1.5 and adjusted p < 0.001), including neutrophils, dendritic cells, Treg, CD8+ T cells, and macrophages (**Fig. 8d**), but not in the ARhigh tumor tissues. Furthermore, by reanalyzing the recently published human PCa scRNA-seq dataset (total patients = 11, total cells = 40,270)^37^, we observed a significant increase in immune cell infiltration within the PCa TME following androgen deprivation therapy (ADT) (**Fig. 8e**). Taken together, these findings suggest that tumor cell-intrinsic AR signaling may play an immunosuppressive effect within the human PCa TME.

**Fig. 8.**
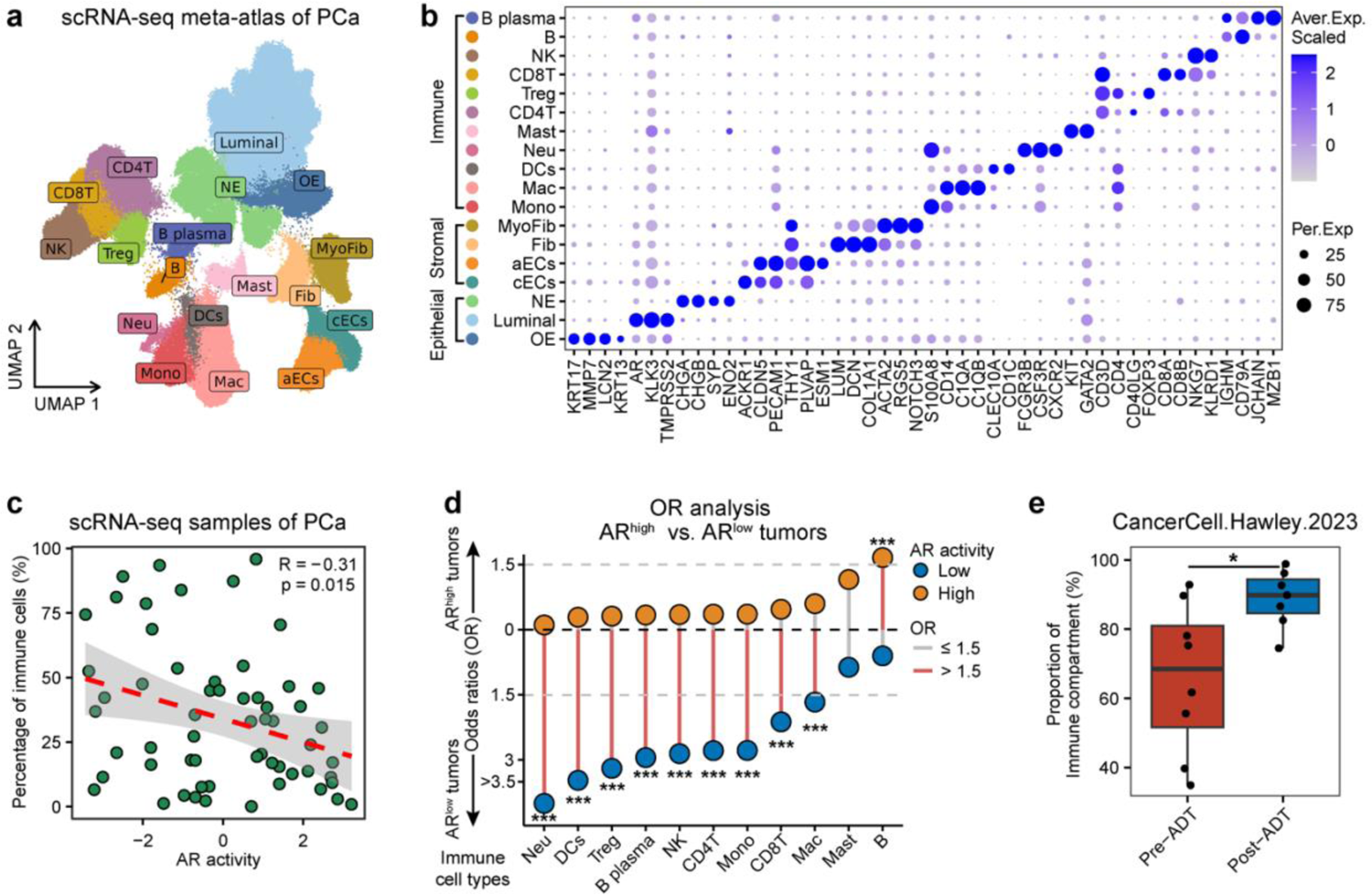
AR activity levels are negatively correlated with immune cell infiltration in TME of human PCa evaluated by scRNA-seq. **a**, UMAP showing the distribution of 18 major cell types in a scRNA-seq atlas of human PCa (tumor samples = 66, cells = 210,879). **b**, Dot plot showing representative marker genes for each cell type. **c**, Pearson correlation analysis between AR activity of epithelial cells and the abundance of immune cells for each sample in the scRNA-seq meta-atlas. **d**, Tissue prevalence of each immune cell type estimated by OR analysis. OR > 1.5 and \*\*\**p* < 0.001 indicates that the cell cluster is preferred to distribute in the corresponding group. **e**, Box plots showing the level of immune cell infiltration between pre-ADT and post-ADT PCa samples from the human PCa scRNA-seq dataset (PCa patients = 11, cells = 40,270) that was generated by Hawley et al. Each dot represents one PCa sample.

## Discussion

Androgens/AR have been implicated as suppressors of immune function in melanoma, prostate and bladder cancer^21, 22, 23, 24, 25^. In this study, we applied a network computational approach to investigate the AR activity profiles in human cancers using 33 bulk tumor RNA-Seq datasets from TCGA cohorts and provide a comprehensive view of its relationship with survival, associated genes, and immune cell infiltration. Our analysis revealed clear negative correlations of pan-cancer AR activity with tumor immune infiltrates in both sexes. We also demonstrated that AR inhibition correlates with enriched immune cell abundance using a unique collection of matched prostate cancer patient biopsies taken before treatment with enzalutamide and upon progression. Notably, our findings suggested that low AR activity, likely reflecting high immune infiltrates and high IFN-gamma immune signature scores in the TME, as associated with improved response to ICB treatment in various cancers.

The role of AR signaling in cancer development is complex and multifaceted. AR can also be activated in a ligand-independent fashion^38^ and signal through interaction with other proteins in the absence of AR binding to DNA, which occurs relatively often in cancer^39^. AR signaling has been shown to interact with several oncogenic signaling pathways involved in promoting growth and resistance across various tumor types, such as AKT/mTOR/PI3K, EGFR, HER2/Neu, and Wnt^39, 40, 41, 42, 43^. Nevertheless, studies correlating AR expression with clinical outcomes have not been consistent. Our analysis showed that PRAD has the highest levels of AR activity compared to other tumors, and higher AR activity is associated with better survival in KIRC. A previous pan-cancer report also indicated that PRAD has the highest AR-score and AR protein expression in TCGA cancers, and high *AR* mRNA and protein levels are associated with better survival in KIRC^44^. Some studies using different cohorts reported similar conclusions, suggesting that AR protein expression is significantly associated with low stage, well-differentiated tumors, and a favorable prognosis in patients with clear cell renal cell carcinoma^45, 46, 47^, yet two studies found that increased expression of *AR* mRNA was associated with poor prognosis^48, 49^. On the other hand, our results showed that a high level of AR activity is associated with poor survival in patients with UCEC and SKCM. UCEC is a hormone-dependent cancer, and a previous study demonstrated that a significant proportion of metastatic endometrial cancers express AR^50^. Additionally, AR is more frequently expressed in metastatic lesions than estrogen and progesterone receptors (ERα and PR), indicating that AR could be a potential therapeutic target, particularly in patients with a high AR to ERα ratio^50^. A previous study found that melanoma cells rely on sustained AR signaling for proliferation and tumorigenesis, and down-modulation correlated with enhanced immune cell infiltration intratumorally in both sexes^23^, which is consistent with the findings in the current study.

It is well-known that males are more susceptible to several non-reproductive malignant cancers^51^. Sex hormones are presumably thought to be the leading actors in sex differences in cancer, as they are known to influence the development and function of the immune system^18^. However, the interaction between sex hormones and the immune system is highly complex^17^, and their specific role in anticancer immunity remains largely unknown. In this study, we observed that AR activity was higher in male tumors in LUSC, while it was higher in female tumors in SARC. Interestingly, the AR activity levels are opposite to the immune infiltration TIMER scores. Notably, nuclear AR has been detected in 70–98% of nonsquamous salivary duct carcinoma (SDC) cases^52^, with a higher prevalence observed in men^53^. Among SDC patients, those who received first-line ADT showed a better response rate and comparable survival outcomes compared to those who underwent first-line chemotherapy^54^. It has also been reported that AR-mediated T-cell exhaustion was more pronounced in male T cells than in female T cells in melanoma^25^ and bladder cancer^24^. Our results, however, demonstrate that AR activity and immune cell abundances are inversely correlated in both sexes. Additionally, AR activity negatively correlated with genes involved in active immune system processes, suggesting the immune suppressive effects of AR in the TME. Herein, we also leveraged a unique dataset with matched biopsies before and after enzalutamide treatment and identified enriched immune infiltrates and IFN-γ signaling activity in the patients’ progression tumor samples that showed AR deactivation.

Another key finding of this study is that the AR activity is highly associated with IFN-γ pathway activity. IFN-γ is a pleiotropic cytokine produced by activated T cells and natural killer (NK) cells within the TME, and it plays a critical role in anti-tumor immunity^55^. Existing evidence has suggested that tumors with a higher expression of genes associated with IFN-γ signaling have a better response to ICB, such as anti-CTLA4 and anti-PD1/PD-L1^30, 56, 57, 58^. We applied ssGSEA to calculate signature scores of three existing IFN-γ-related signatures to represent IFN-γ signaling activity in tumor samples. More specifically, the Hallmark IFN gamma response pathway and the T cell-inflamed GEP signature contain IFN-γ-responsive genes associated with antigen presentation, chemokine expression, cytotoxic activity, and adaptive immune resistance^29, 30^. The ICR 20-gene signature reflects the strength of cytotoxic response characterized by T helper type-1 signaling, chemoattraction, cytotoxic function, and immune checkpoint-related genes^31, 58^. Our results from both TCGA and GTEx demonstrated a significant negative correlation between AR activity and the three IFN-γ-associated signature scores in both tumor and normal tissues, suggesting AR has a negative impact on anti-tumor immune response. The data imply that deactivation of AR in the TME reflects higher IFN-γ signaling activity, which might be an indicator of responsiveness to ICB therapy. Indeed, we analyzed the transcriptome of baseline tumor biopsies from six independent datasets of multiple cancer types treated with anti-PD1/PD-L1 or anti-CTLA-4 and identified non-responders had higher AR activity than the responders. Our findings are in line with a recent publication in *Nature*; Guan et al. reported preclinical and clinical evidence elucidating AR as a negative modulator of CD8^+^ T cells response to anti-PD1/PD-L1 treatment^22^.

There are some limitations to this study. There are mixed reports on the impact of high AR activity and survival outcomes. Therefore, it is important to further explore the underlying mechanisms between AR signaling and anti-cancer immunity, as well as its correlations with clinical outcomes across cancer contexts. In addition, we mainly employed the TCGA database to perform these analyses, in which primary tumors are the majority of samples, and the included studies with ICB treatments did not cover all cancer types. Further investigation of our hypothesis using a large sample size with different cancer types, including subtypes of each cancer, is necessary to confirm the role of AR activity in predicting ICB response across all cancer types. Rigorous pan-cancer studies are required in the future to offer additional independent validation.

In summary, we applied regulon-based bioinformatics approaches and determined that AR activity is strongly correlated with tumor immune infiltration and ICB response across multiple tumor lineages. We conclude that AR activity is highly correlated with specific immune responses and infiltrates, suggesting that combining immunotherapies with AR blockade may be a promising treatment approach for several cancers, particularly in melanoma, in which we demonstrated that AR activity is a distinctive risk factor for survival and has a strong negative correlation with immune infiltrates.

## Methods

### Data source and processing

Bulk RNA sequencing (RNA-seq) data and corresponding clinical data for 33 cancer types from The Cancer Genome Atlas (TCGA) consortium were retrieved from The National Cancer Institute (NCI) Genomic Data Commons (GDC) (https://portal.gdc.cancer.gov/) via TCGAbiolinks R/ Bioconductor package (version 2.24.1)^59, 60, 61^. The Genotype-Tissue Expression (GTEx)^62^ data were downloaded from the portal (https://www.gtexportal.org/home/downloads/adult-gtex#bulk_tissue_expression). The independent datasets of clinical trials were accessed from Gene Expression Omnibus (GEO, https://www.ncbi.nlm.nih.gov/ geo/) database (GSE145996^63^, GSE135222^64^, GSE126044^65^), the database of Genotypes and Phenotypes (dbGaP, https://www.ncbi.nlm.nih.gov/gap/; phs000452.v2.p1^66^), or publicly available source data (Yang et al.^67^, Westbrook et al.^68^, and Guan et al.^22^). R package “edgeR” (version 3.38.1)^69, 70, 71^ was utilized to process the downloaded bulk RNA-seq data value. The downloaded RNA-seq data were then converted into transcripts per kilobase million (TPM) values and further transformed as log(1+TPM) for downstream analyses. Three independent single-cell RNA-seq datasets were collected from previous publication (Zhang et al.^35^, Zheng et al.^36^, and Hawley et al.^37^).

### Estimating single-sample TF activity levels

To measure AR activity and the activity of estrogen receptors, progesterone, and glucocorticoid receptors of each sample from datasets, single-sample Virtual Inference of Protein-activity by Enriched Regulon analysis (VIPER) was performed using the VIPER R package (version 1.30.0)^26^. A log1p transformed TPM gene expression matrix and a regulatory network were used as inputs for VIPER analysis. The TF regulons (the regulatory network) used in this study was curated from several databases as previously described^72^.

### Survival analysis

Survival analysis was performed using univariate Cox proportional hazard models to evaluate the association between AR activity levels and clinical outcomes in each cancer type. *P* values less than 0.05 were considered statistically significant. To ensure proper use of the clinical outcome endpoints in our pan-cancer study, we selected overall survival (OS) or progression-free interval (PFI) endpoints for each disease type as the outcome endpoints according to the provided recommendations from a published article (Liu et al., Table3)^27^, which demonstrated a standardized dataset named the TCGA Pan-Cancer Clinical Data Resource (TCGA-CDR). Among 33 TCGA cancer types, PFI was used in BRCA, DLBC, LGG, TGCT, THYM, PRAD, READ, and THCA, while OS was used in other cancer types.

### Identification of AR activity-correlated genes and over-representation analysis

To identify genes commonly correlated with AR activity in 33 TCGA cancer types, we performed Pearson correlation analysis to assess the correlations between AR activity and each gene expression in each cancer type. We then overlapped the significantly correlated gene lists in the 33 cancer types to identify the 31 genes common in all 33 TCGA cancer types.

The “ClusterProfiler” R package (version 4.4.1) ^73, 74^ was used to perform GO and Reactome over-representation analysis for the exploration of pathways and biological processes relating to the 31 AR-activity correlated genes. The biological processes within the GO domain were investigated. Enrichment pathways and categories at adjusted p value < 0.05 are selected as statistically significant and the top 15 significant enrichment GO BP or Reactome categories are visualized in the dot plots.

### Estimation of tumor-infiltrating immune cells

Tumor Immune Estimation Resource 2.0^75, 76, 77^ (TIMER2.0; http://timer.cistrome.org/) web server is a comprehensive resource for evaluating the abundances of tumor-infiltrating immune cells across diverse cancer types. We used it to obtain the infiltration levels of B cells, CD4^+^ T cells, CD8^+^ T cells, Neutrophil, Macrophage, and Myeloid dendritic cells in 32 cancer types from TCGA database estimated by TIMER immune deconvolution algorithm. We then explored the association between AR activity and immune infiltration using Pearson correlation analysis.

### Calculating IFN-γ related gene expression signature and TLS signature scores

We collected three published IFN-γ-related gene expression signatures and one TLS signature associated with prognostic value to immunotherapy, including Hallmark interferon gamma response gene set (IFNG MSigDB hallmark gene set^29^), 18-gene T cell-inflamed GEP^30^, the immunologic constant of rejection (ICR) 20-gene signature^31^ and the TLS signature (Cabrita et al.^32^). Gene lists of each signature are in Supplementary Table 3. To estimate the activity of these gene signature scores, we used the single-sample gene-set enrichment analysis (ssGSEA) implemented in GSVA R package (Version 1.44.1)^78^ to calculate sample-wise absolute enrichment scores of these signatures in each sample. The ssGSEA algorithm is a rank-based method to assess the expression levels of genes of a gene signature against all other genes in each sample within a given dataset. Log transformed gene expression profiles and gene sets from the published studies were used as input to ssGSEA.

### Correlation analysis

We computed the correlation between two continuous variables using Pearson’s correlation coefficients. A significance threshold of *p* < 0.05 (Pearson’s correlation test) was applied to determine the significance of correlation. When calculating the correlation between immune pathways and AR activities, the overlap genes between the gene signatures and the AR regulon were removed from the pathway gene sets.

### Master regulator analysis and pathway analysis

RNA-seq data of 21 matched tumor biopsy samples of prior to enzalutamide and at the time of progression (Baseline and Progression)^68^ were used to evaluate differential TF activities and to perform IFN-γ signatures, immune subpopulations, and GO biological process gene-set enrichment analysis (GSEA)^79^. Differential gene expression analysis was first performed using DESeq2 (version 1.32.0)^80^. Gene expression differences were considered significant when adjusted *p*-value < 0.05. TF activity was inferred by msVIPER algorithms provided in the VIPER R package (version 1.26.0)^26^. The Wald test statistic results from DESeq2 output were served as a gene list input data for VIPER analysis. The transcriptional regulatory network used in this study was curated from four databases as previously described^72^. Pathway analysis was performed using GSEA function implemented in the “ClusterProfiler” R package (version 4.4.1)^73, 74^. The Wald test statistic results from DESeq2 output were served as a pre-ranked gene list input of GSEA. The gene sets were considered to be activated if their adjusted-p value was less than 0.05 and the NES (Normalized Enrichment Score) was greater than 0.

### Single-cell analysis methodology

The recently published human PCa scRNA-seq meta-atlas^35^ and the scRNA-seq dataset^37^ were reanalyzed using the Seurat pipeline (v4.1.0). Briefly, the BBKNN algorithm^81^ was applied to mitigate potential batch effects. Uniform Manifold Approximation and Projection (UMAP) with the Leiden algorithm was employed for clustering and visualizing single-cell distribution. Cell clusters were annotated based on previously reported cell type marker genes of human PCa^35^ and the combined automatic annotation method Celltypist^82^. Odds ratios (OR) were calculated to quantify the tissue preference of each immune cell type in PCa tissues with different AR activity statuses^36^. OR > 1.5 indicates that the cell type is preferred to distribute in the corresponding tissue^36^.

### Data analysis of the independent ICB datasets

To further investigate the clinical utility of single-sample AR activity in different cancer types, we obtained five independent datasets from published studies and one of our studies with bulk RNA-Seq data for tumor samples collected from patients with clinical information about immune checkpoint inhibitor therapeutic responses. We used baseline gene expression profiles for our analyses. The GSE145996^63^ and phs000452.v2.p1^66^ datasets included 33 non-responders and 23 responders of melanoma patients. The GSE135222^64^ and GSE126044^65^ datasets included 30 non-responders and 13 responders of non-small-cell lung cancer (NSCLC) patients. Yang et al.^67^ dataset included tumor samples collected from The Investigator-initiated Phase 2 Study of Pembrolizumab Immunological Response Evaluation (INSPIRE) cohorts of advanced solid cancer patients treated with pembrolizumab, which included Neck Squamous Cell Cancer (HNSCC), triple-negative breast (TNBC), high-grade serous carcinoma (HGSC), melanoma, and other mixed solid tumors (MST). Using the classifications in Yang’s study, 18 pembrolizumab-high sensitivity/clinical benefit (the HS/CB group; responder) and 22 low sensitivity (the LS group; non-responder) samples were analyzed in our study.

### Statistical analysis

All data processing, statistical analyses, and plotting were conducted in the R statistical computing environment software v4.2.0 (https://www.R-project.org/). Cox regression analyses were performed via the survival R package (version 3.3-1)^83, 84^. We calculated the correlation between two continuous variables using the Pearson’s correlation coefficients. The threshold of *p* < 0.05 (Pearson’s correlation test) indicates the significance of correlation. Wilcoxon rank-sum tests were performed on continuous measures between two groups to examine differences in distributions. All statistical tests were two-sided, and a *p*-value < 0.05 was considered as statistically significant.

## Data availability

The publicly available datasets used in this study can be accessed from multiple sources. The datasets from Genomic Data Commons (GDC) (https://portal.gdc.cancer.gov/) are available via TCGAbiolinks R/Bioconductor package (version 2.24.1)^59, 60, 61^, while the Gene Expression Omnibus (GEO, https://www.ncbi.nlm.nih.gov/geo/) database contains GSE145996^63^, GSE135222^64^, GSE126044^65^, and dbGaP (https://www.ncbi.nlm.nih.gov/gap/; phs000452.v2.p1^66^) is another source. Additionally, the source data from literatures such as Zhang et al.^35^, Zheng et al.^36^, Hawley et al.^37^, Yang et al.^67^, Westbrook et al.^68^, and Guan et al.^22^ were used in this study. Any additional data are either available within the Article or in the Supplementary Information files. The author can provide any remaining data upon reasonable request.

## Code availability

All packages utilized in this study are publicly accessible. While this study does not generate any original code, the corresponding author is available to provide further details upon request.

## Acknowledgements

This work was supported by the following funding: Department of Defense Idea Development Award W81XWH2110539, NIH R01GM147365 and Silver Family Innovation Foundation Award (to Z.X.); NIH R37CA263592 (to A.E.M.). Breast Cancer Research Foundation and NIH U01CA253472 and U01CA217842 (to G.B.M.). NIH R01CA251245, NIH R01CA282005, NCCN/Pfizer/Astellas Award, and Joint Institute for Cancer Research Award (to J.J.A.). Y.M.H. was supported by a postdoctoral fellowship of Portland Oral Health Research Training (PORT) program NIH T90 DE030859 (to H. W.). The content is solely the responsibility of the authors and does not necessarily represent the official views of the funders.

## Author contributions

Z.X. conceived the idea. Z.X., A.E.M. and Y.M.H. contributed to the study design. Y.M.H., F.Z., and Z.X. performed computational analyses. Y.M.H., F.Z., Z.X., A.E.M., J.J.A., and J.N.G. interpreted data. C.C. and X.Z. contributed to the data analysis. G.B.M., A.K., G.V.T., and H.W. contribute to the analytic strategies insights. Z.X. supervised the study. Y.M.H. and Z.X. wrote the manuscript. All other authors provided critical feedback and approved the final manuscript.

## Competing interests

G.B.M. is SAB/Consultant for AstraZeneca, BlueDot, Chrysallis Biotechnology, Ellipses Pharma, ImmunoMET, Infinity, Ionis, Lilly, Medacorp, Nanostring, PDX Pharmaceuticals, Signalchem Lifesciences, Tarveda, Turbine and Zentalis Pharmaceuticals; stock/ options/financial: Catena Pharmaceuticals, ImmunoMet, SignalChem, Tarveda and Turbine; licenced technology: HRD assay to Myriad Genetics, and DSP patents with NanoString. A.E.M. received research funding from AstraZeneca. J.J.A. has received consulting income from Fibrogen, Astellas, and Bristol Myers Squib. and research support to his institution from Beactica, a Pfizer/Astellas/NCCN research award, and Zenith Epigenetics. The remaining authors declare no competing interests.

## Materials & Correspondence

Correspondence and requests for materials should be addressed to Z.X.

