## Supplementary Figures for "Androgen receptor activity inversely correlates with immune cell infiltration and immunotherapy response across multiple cancer lineages"

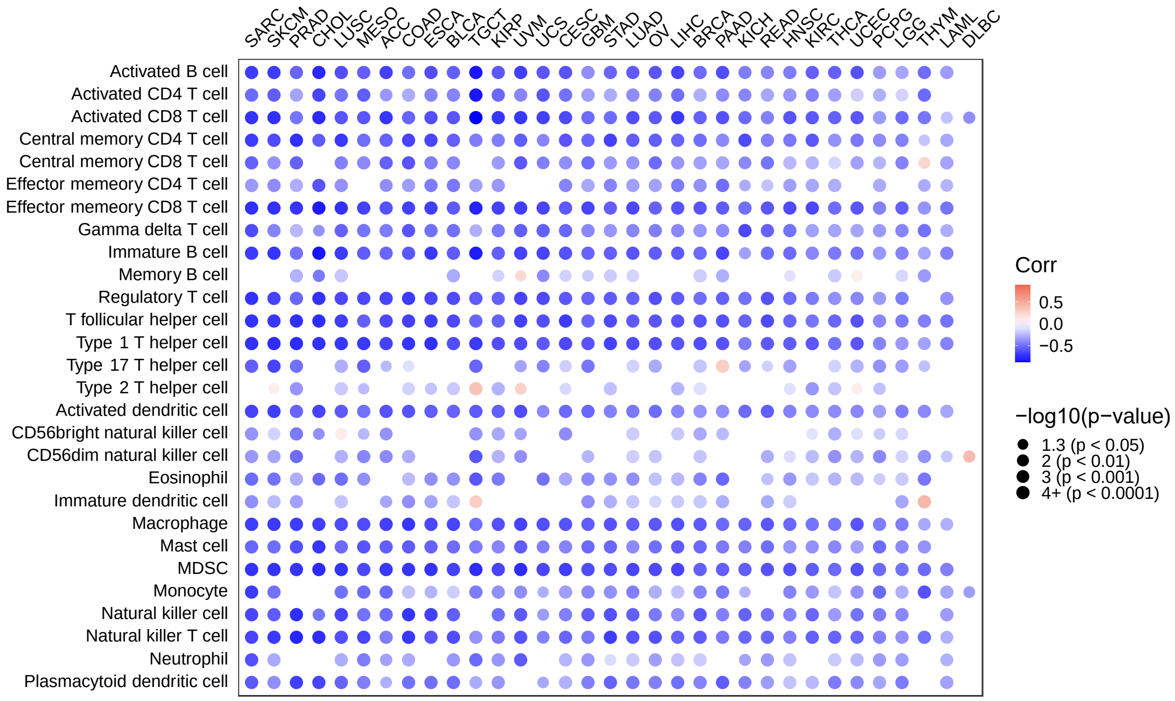


**Supplementary Figure 1:** Correlation between AR activity and ssGSEA enrichment scores of 28 immune cell types across 33 cancer types. The p-value shown was determined using a two-tailed Pearson correlation. Positive correlation coefficients are displayed in orange, and negative correlation coefficients in blue. The color intensity is proportional to the correlation coefficients. The circle size corresponds to the *p*-value, while correlation coefficients with $p > 0.05$ are left blank. ssGSEA: single-sample Gene Set Enrichment Analysis.

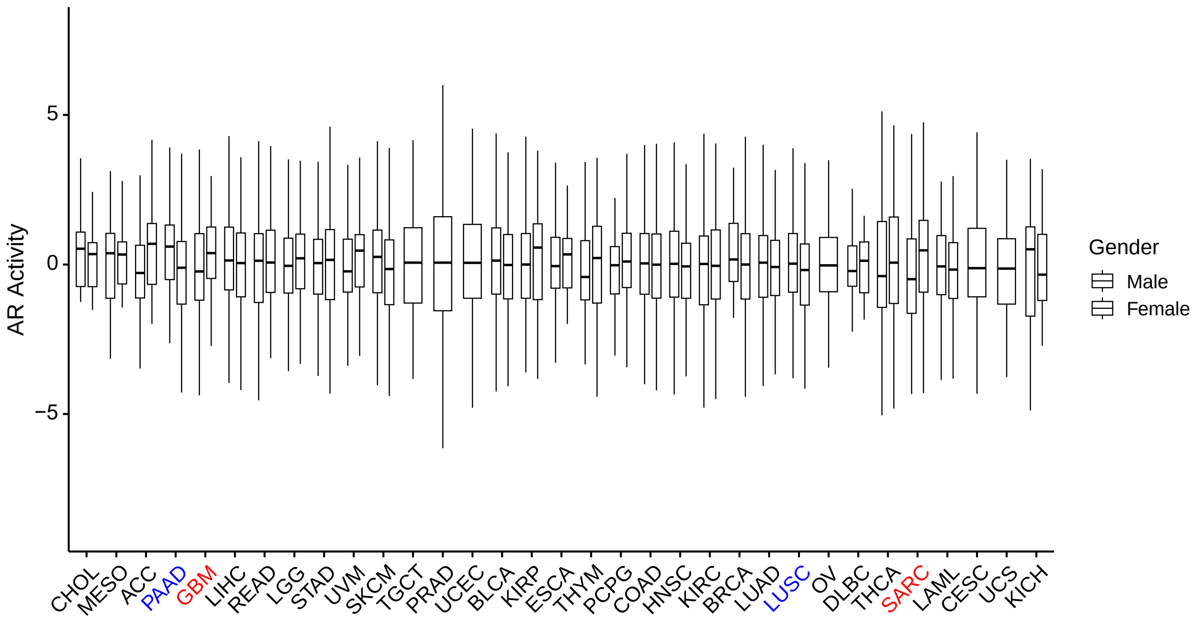


**Supplementary Figure 2:** Boxplots illustrating AR activity between male (blue) and female (pink) patients in 33 TCGA cancer cohorts. Blue texts represent AR activity is significantly higher in male patients than in females, while pink texts represent AR activity is significantly higher in female patients than in males. Statistical significances were calculated by two-sided Wilcoxon rank-sum tests between males and females. The center line indicates median, bounds of the box indicate upper and lower quartiles, whiskers indicate minimum and maximum, and outliers are marked with dots.

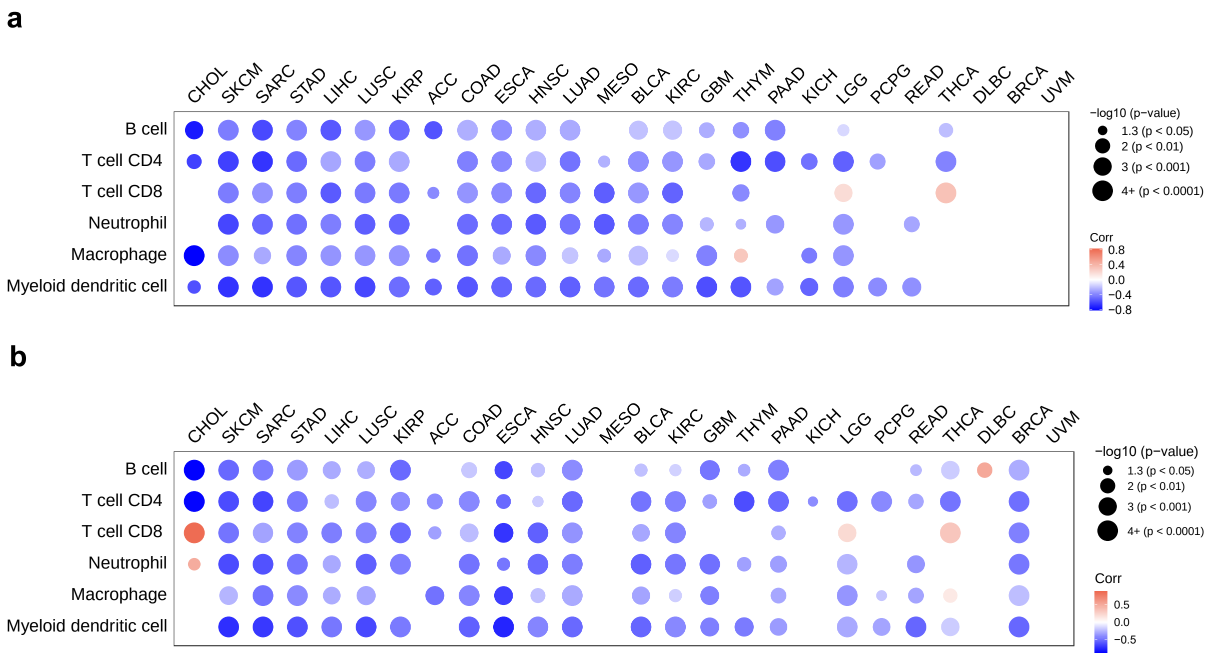


**Supplementary Figure 3.** Correlations between AR activity and six immune cell populations (B cells, CD4$^{+}$ T cells, CD8$^{+}$ T cells, macrophages, dendritic cells, and neutrophils) in men (**a**) and women (**b**) across 26 cancer types. Each circle represents a correlation coefficient value analyzed by two-tailed Pearson correlation test. Positive correlation coefficients are displayed in orange color and negative correlation coefficients in blue color. The color intensity is proportional to the correlation coefficients. The circle size is proportional to the *p*-value, while correlation coefficients with $p > 0.05$ are blank. Cancer types (column) are sorted in order of decreasing mean of correlation coefficient. The enrichment scores of immune infiltration levels are determined by TIMER algorithm.

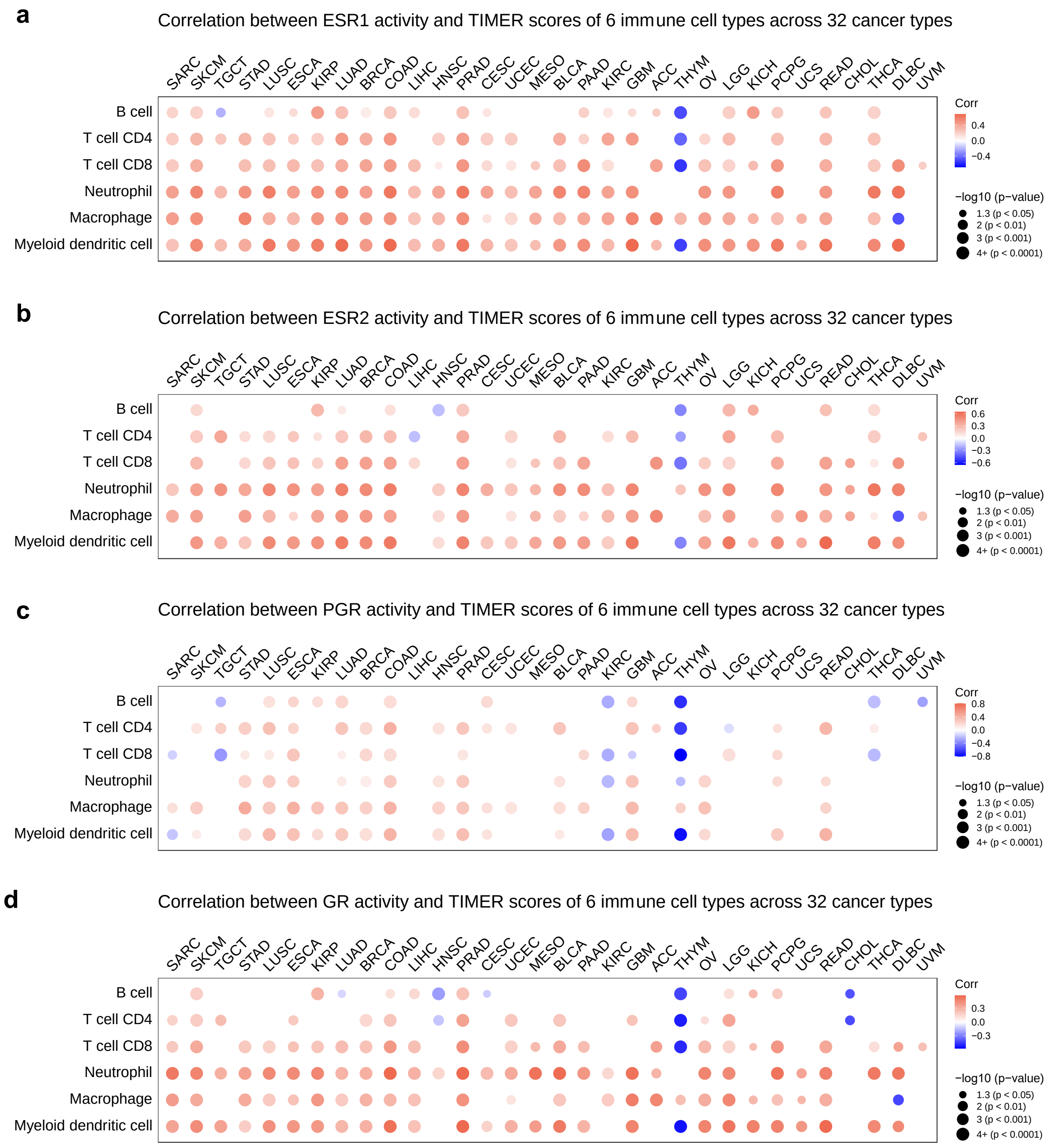


**Supplementary Figure 4**

**Supplementary Figure 4:** Dot plots illustrate the correlations between **a** estrogen receptor 1 (ESR1), **b** estrogen receptor 2 (ESR2), **c** progesterone receptor (PGR), and **d** glucocorticoid receptor (GR) activity, and six immune cell populations (B cells, $CD4^+$ T cells, $CD8^+$ T cells, macrophages, dendritic cells, and neutrophils) across 32 cancer types in TCGA cohorts (LAML is not applicable in TIMER calculation). Each circle represents a correlation coefficient value analyzed using a two-tailed Pearson correlation test. Positive correlation coefficients are displayed in orange, and negative correlation coefficients are displayed in blue. The color intensity is proportional to the correlation coefficients. The circle size is proportional to the *p*-value, while correlation coefficients with $p > 0.05$ are left blank. The cancer types in the column are arranged in the same order as illustrated in Fig. 3a. The enrichment scores of immune infiltration levels are determined using TIMER algorithm.

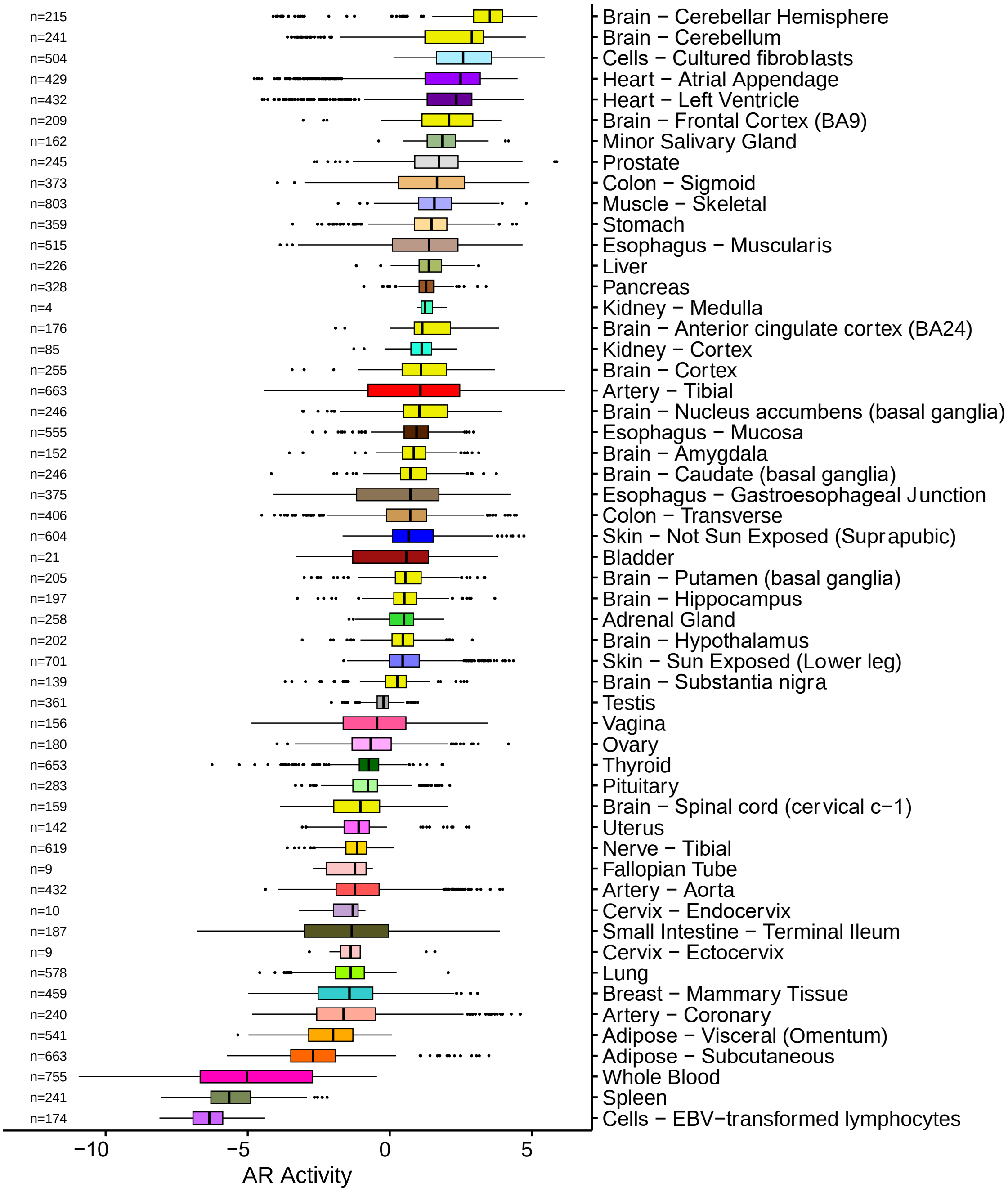


**Supplementary Figure 5:** Boxplots displaying AR activity sorted in order of decreasing AR activity median among 54 GTEx tissue types. Values on the left correspond to the total number of tissue samples (n) analyzed in each tissue cohort. The center line indicates median, bounds of the box indicate upper and lower quartiles, whiskers indicate minimum and maximum, and outliers are marked with dots. GTEx: The Genotype-Tissue Expression.

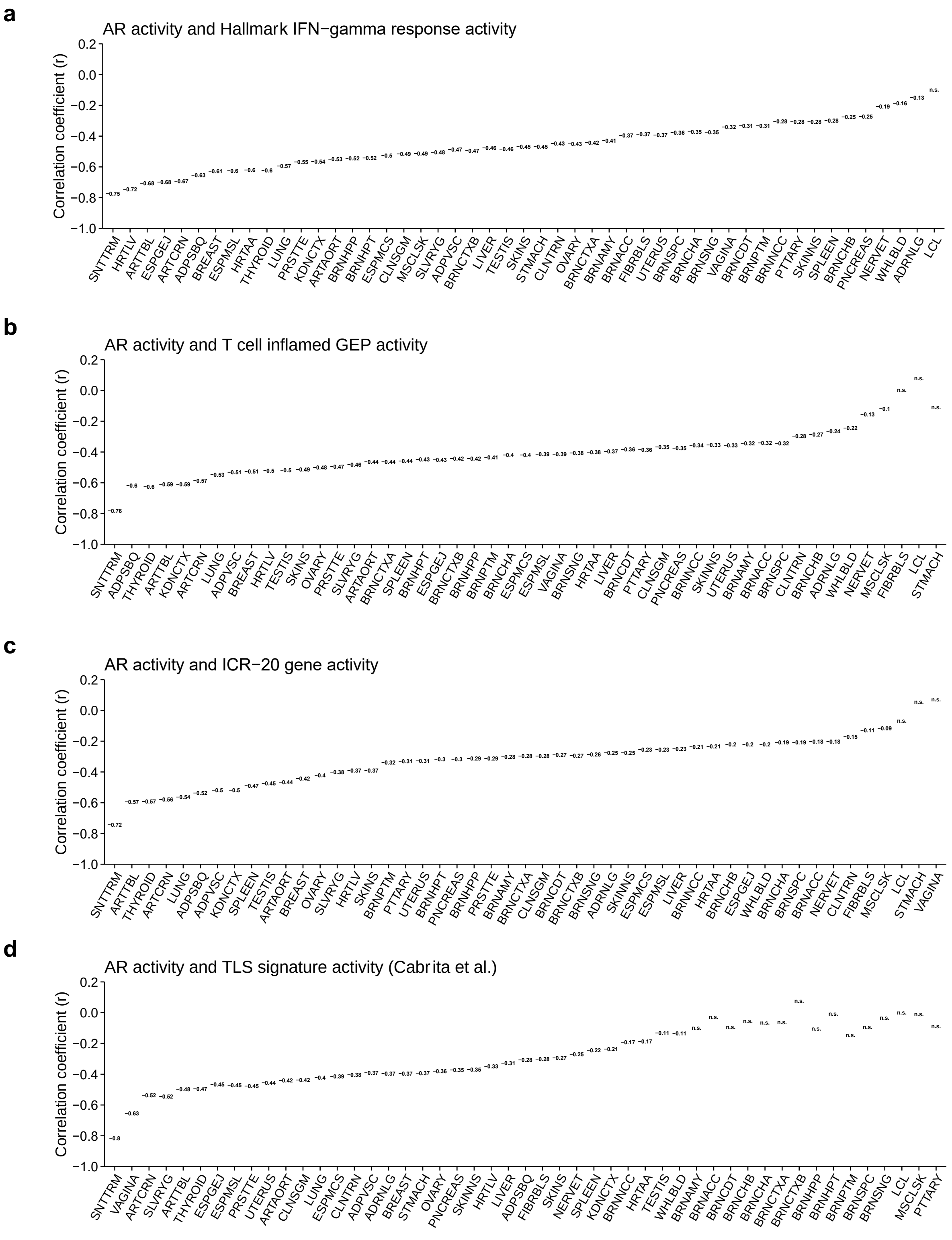


**Supplementary Figure 6**

**Supplementary Figure 6:** Bar plots (**a, b, c, d**) display the Pearson correlation of AR activity with four parameters: **a**) Hallmark IFN-γ pathway, **b**) T cell–inflamed GEP, **c**) ICR 20-gene, and **d**) TLS signature activity scores within each tissue type from the GTEx database. Tissue types with fewer than 30 samples were excluded from the analysis. The tissue type abbreviations and sample numbers are listed in Supplementary Table 2. The X-axis represents tissue types arranged in descending order of correlation coefficient values (Y-axis). The text below each bar indicates the Pearson correlation coefficients. Bars on the far right with a p-value greater than 0.05 are colored in light grey. Definitions: GEP: Gene Expression Profile, ICR: Immunologic Constant of Rejection, TLS: Tertiary Lymphoid Structures, GTEx: Genotype-Tissue Expression. 'ns' indicates non-significant correlations.

**Supplementary Table 1. TCGA Study Abbreviations and Datasets Evaluated**

| Symbol | Name | Tumor Samples (n) |
|---|---|---|
| ACC | Adrenocortical carcinoma | 79 |
| BLCA | Bladder urothelial carcinoma | 414 |
| BRCA | Breast invasive carcinoma | 1108 |
| CESC | Cervical squamous cell carcinoma and endocervical adenocarcinoma | 306 |
| CHOL | Cholangiocarcinoma | 36 |
| COAD | Colon adenocarcinoma | 478 |
| DLBC | Lymphoid neoplasm diffuse large B-cell lymphoma | 48 |
| ESCA | Esophageal carcinoma | 162 |
| GBM | Glioblastoma multiforme | 168 |
| HNSC | Head and neck squamous cell carcinoma | 502 |
| KICH | Kidney chromophobe | 65 |
| KIRC | Kidney renal clear cell carcinoma | 539 |
| KIRP | Kidney renal papillary cell carcinoma | 289 |
| LAML | Acute myeloid leukemia | 151 |
| LGG | Brain lower grade glioma | 528 |
| LIHC | Liver hepatocellular carcinoma | 374 |
| LUAD | Lung adenocarcinoma | 535 |
| LUSC | Lung squamous cell carcinoma | 502 |
| MESO | Mesothelioma | 86 |
| OV | Ovarian serous cystadenocarcinoma | 379 |
| PAAD | Pancreatic adenocarcinoma | 178 |
| PCPG | Pheochromocytoma and paraganglioma | 183 |
| PRAD | Prostate adenocarcinoma | 499 |
| READ | Rectum adenocarcinoma | 166 |
| SARC | Sarcoma | 263 |
| SKCM | Skin cutaneous melanoma | 471 |
| STAD | Stomach adenocarcinoma | 375 |
| TGCT | Testicular germ cell tumors | 139 |
| THCA | Thyroid carcinoma | 510 |
| THYM | Thymoma | 119 |
| USEC | Uterine corpus endometrial carcinoma | 552 |
| UCS | Uterine carcinosarcoma | 56 |
| UVM | Uveal melanoma | 80 |

**Supplementary Table 2. GTEx Tissue Type Abbreviations and Sample Numbers**

| Abbreviation | Tissue Site Detail | Sample number (n) |
|---|---|---|
| ADPSBQ | Adipose - Subcutaneous | 663 |
| ADPVSC | Adipose - Visceral (Omentum) | 541 |
| ADRNLG | Adrenal Gland | 258 |
| ARTAORT | Artery - Aorta | 432 |
| ARTCRN | Artery - Coronary | 240 |
| ARTTBL | Artery - Tibial | 663 |
| BLDDER | Bladder | 21 |
| BRNAMY | Brain - Amygdala | 152 |
| BRNACC | Brain - Anterior cingulate cortex (BA24) | 176 |
| BRNCDT | Brain - Caudate (basal ganglia) | 246 |
| BRNCHB | Brain - Cerebellar Hemisphere [Frozen] | 215 |
| BRNCHA | Brain - Cerebellum [PAXgene] | 241 |
| BRNCTXA | Brain - Cortex [PAXgene] | 255 |
| BRNCTXB | Brain - Frontal Cortex (BA9) [Frozen] | 209 |
| BRNHPP | Brain - Hippocampus | 197 |
| BRNHPT | Brain - Hypothalamus | 202 |
| BRNNCC | Brain - Nucleus accumbens (basal ganglia) | 246 |
| BRNPTM | Brain - Putamen (basal ganglia) | 205 |
| BRNSPC | Brain - Spinal cord (cervical c-1) | 159 |
| BRNSNG | Brain - Substantia nigra | 139 |
| BREAST | Breast - Mammary Tissue | 459 |
| FIBRBLS | Cells - Cultured fibroblasts | 504 |
| LCL | Cells - EBV-transformed lymphocytes | 174 |
| CVXECT | Cervix - Ectocervix | 9 |
| CVSEND | Cervix - Endocervix | 10 |
| CLNSGM | Colon - Sigmoid | 373 |
| CLNTRN | Colon - Transverse | 406 |
| ESPGEJ | Esophagus - Gastroesophageal Junction | 375 |
| ESPMCS | Esophagus - Mucosa | 555 |
| ESPMSL | Esophagus - Muscularis | 515 |
| FLLPNT | Fallopian Tube | 9 |
| HRTAA | Heart - Atrial Appendage | 429 |
| HRTLV | Heart - Left Ventricle | 432 |
| KDNCTX | Kidney - Cortex | 85 |
| KDNMDL | Kidney - Medulla | 4 |

**Supplementary Table 2. GTEx Tissue Type Abbreviations and Sample Numbers (Continued)**

| Abbreviation | Tissue Site Detail | Sample number (n) |
|---|---|---|
| LIVER | Liver | 226 |
| LUNG | Lung | 578 |
| SLVRYG | Minor Salivary Gland | 162 |
| MSCLSK | Muscle - Skeletal | 803 |
| NERVET | Nerve - Tibial | 619 |
| OVARY | Ovary | 180 |
| PNCREAS | Pancreas | 328 |
| PTTARY | Pituitary | 283 |
| PRSTTE | Prostate | 245 |
| SKINNS | Skin - Not Sun Exposed (Suprapubic) | 604 |
| SKINS | Skin - Sun Exposed (Lower leg) | 701 |
| SNTTRM | Small Intestine - Terminal Ileum | 187 |
| SPLEEN | Spleen | 241 |
| STMACH | Stomach | 359 |
| TESTIS | Testis | 361 |
| THYROID | Thyroid | 653 |
| UTERUS | Uterus | 142 |
| VAGINA | Vagina | 156 |
| WHLBLD | Whole Blood | 755 |
